# ANTIMICROBIAL RESISTANCE AND VIRULENCE GENES PROFILING OF *PROTEUS* SPECIES FROM POULTRY FARMS IN LAFIA, NIGERIA

**DOI:** 10.1101/2021.01.07.425673

**Authors:** Mojisola Christiana Owoseni, Odula Oyigye, Bashiru Sani, Jebes Lamin, Amara Chere

## Abstract

The poultry industry is important in boosting food sustainability of a population; however, the poultry environment and products are susceptible to pathogen contamination arising from poor farm hygienic conditions. This study investigated the prevalence, antimicrobial resistance and virulence profile of *Proteus* species from the environment and products of four selected poultry farms in Lafia, Nasarawa State. Farm samples (n =216) comprising feed (64), drinking water (64) and swabs from eggshells (88) were collected and analysed for detection of *Proteus* species using cultural, biochemical and microscopic techniques. Antibiotics susceptibilities of isolates were also determined, and virulence genes were confirmed using Polymerase Chain Reaction. Of the total studied samples, 34.26% (74/216) were positive for *Proteus* species. *Proteus* species were more prevalent in drinking water samples (37.84%) and feed samples (33.78%) and least prevalent in eggshells (28.38%). *Proteus* species (n= 74) comprised *P. mirabilis* 78% (58/74) and *Proteus vulgaris* 22% (16/74). *P. mirabilis* was markedly higher than *P. vulgaris* in all the four farms sampled. Farms A and D had the highest prevalence of *Proteus* species, while Farms A and C (80%), and Farm D (25) has the highest prevalence of *P. mirabilis* and *P. vulgaris* respectively. The isolates’ prevalence rate within and between farms, sample type, and species was not statistically significantly different (p≥0.05) from the other farms. Isolates were 100% susceptible to Amikacin and showed the highest resistance (25.7%) to tetracycline. Molecular analysis of the virulence genes of *Proteus* species revealed the presence of *rsbA*, *ureC* and *luxS* virulent genes in all the test isolates. Data generated indicates a high level of multidrug-resistant pathogenic strains of *Proteus* circulating in poultry farms in Lafia, Nigeria, which potentiates a significant risk of transmission of pathogenic *Proteus* via the food chain.

## 1. Introduction

The poultry industry is a major component of the Nigerian economy, providing a source of income to farmers and serving as a leading source of high-quality protein for the fast-growing population due to the affordability and acceptability of their products (Bettridge et al., 2014). However, the incidence of microbial infections from infected birds with clinical diseases in the poultry farm environment and products has threatened public health and food safety and may cripple economic gains derived from this industry (Foti et al., 2011; Menghistu et al., 2011).

*Proteus* poses a significant challenge to both humans and animals worldwide and is prevalent in several food and animals including poultry (Lei et al., 2014). There are now multiple resistant forms of this bacterium which indicates a major food issue (Lei et al., 2016). With several foodborne poisoning attributed to *Proteus* group of bacteria and rising incidence of *Proteus* induced foodborne infections, it is pertinent that control programs and prophylactic measurements be developed to prevent and combat outbreaks of foodborne infections/poisoning from poultry and poultry products (Ram et al., 2019). Poultry and other poultry products are believed to be the primary vehicle for the transmission of *Proteus* (Firildak et al., 2015). Poultry might have *Proteus* strains in their droppings and on their bodies (feathers, feet and beaks), even when they appear healthy and clean. The *Proteus species* can stick to cages, coops, feed and water dishes, hay, plants and soil in the area where the birds live; also eggshells may become contaminated with *Proteus* from poultry droppings (poop) or the area where they are laid (Jambalang et al., 2017). From a public health perspective, the number of eggs and animals affected by *Proteus* is a risk factor for human disease or infection (Awad-Alla et al., 2010).

Human cases of *Proteus* are typically acquired through the consumption of contaminated food. *Proteus* usually spread through the faecal-oral route (contamination of hands or objects with bacteria shed in the stool) (Tonkić et al., 2010). *Proteus* in chicken droppings might be transmitted to vulnerable workers while handling infected chicken directly or through faecal-contaminated poultry products (Barua et al., 2013; Lima-Filho et al., 2013). Generally, there are two possible routes of egg contamination by *Proteus*. Eggs can be contaminated by penetration of the bacterium through the eggshell from the colonised gut or from contaminated faeces during or after oviposition (horizontal transmission). Horizontal transmission occurs following ingestion of food or water already contaminated with faeces of clinically infected birds or carriers, presence of dead chickens, poultry farm attendants and contaminated feeds (Armbruster et al., 2014). The second possible route is by direct contamination of the yolk, albumen, eggshell membranes or eggshells before oviposition, originating from the infection of reproductive organs with *Proteus* (vertical transmission) (Momani et al., 2017; Moosavy et al., 2015).

Consumption of raw/undercooked eggs has consistently been identified as the primary risk factor for *Proteus mirabilis* (MD Salihu, 2015). *Proteus* is known to cause human urinary tract infection (UTI), nosocomial infection, wound infection (Ebringer and Rashid, 2014; Jacobsen et al., 2008) and showed clear history of zoonosis in vast host range with the emergence of multidrug resistance (MDR) in recent years (Tonkić et al., 2010). Multidrug-resistant *Proteus* may be transmitted among poultry farm workers who may transmit the pathogen in the surrounding environment.

Humans also use many classes of antimicrobial agents used in animals, and there is a potential selection and spread of antimicrobial-resistant bacteria or genes from animals to humans through the food chain (Food and Agriculture Organization (FAO), 2016). The indiscriminate use of antibiotics or congeners has created enormous pressure for the selection of antimicrobial resistance among bacterial pathogens worldwide, including *Proteus* strains found in poultry products and poultry environment (Nahar et al., 2014).

The virulence of the *Proteus* species is caused by several factors which are regulated and expressed by virulence genes encoded in operons (Kumar Trivedi, 2015; Pathirana et al., 2018). These virulence genes increase the pathogenicity of *Proteus* species among which include urease which is the most important enzyme for kidney and bladder stone formation (Mohamed et al., 2014) and enables it to produce an environment in which it can survive (Aboh et al., 2015). The *luxS* gene is involved in the synthesis of autoinducer 2 (AI-2) secreted by bacteria and used to communicate both the cell density and the metabolic potential of the environment (Badi and Sepahi, 2014). Swarming behaviour of *P. mirabilis* mediated by *rsbA* gene has been associated with biofilm formation and extracellular polysaccharide formation (Różalski et al., 2012). Continuous monitoring and surveillance of poultry farms is pertinent to establish possible transmission routes of microbial pathogens from poultry farms via the food chain.

This study investigated the prevalence, antimicrobial resistance and virulence gene profiles of *Proteus* species isolated from Poultry farm samples in Lafia, Nigeria.

## 2. Material and Methods

### 2.1 Sample collection and bacterial isolation

A cross-sectional study was conducted for two months between May and June 2019, and a total number of 216 samples from four commercial poultry farms in Lafia, Nasarawa State were collected. The sample size was determined by using the prevalence of 16.67% reported by (Esther Chat et al., 2019) and the following the equation described by (Naing et al., 2006) as

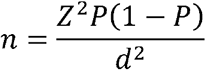

Where: n = sample size

P = prevalence from a previous study = 16.67% = 0.17.

Z = standard normal distribution at 95% confidence interval= 1.96.

D = absolute desired precision at 5% = 0.05.

Samples included poultry feed (n = 64), drinking water (n=64) and unbroken eggshells (n = 88) from four selected poultry farms in equal amounts during early morning hours on a weekly interval. Swab samples collected from surfaces of eggshells were inoculated into buffered peptone water (BPW) (Oxoid, UK) in screw-capped bottles, incubated at 37 °C for 24 h and thereafter pre-enriched in Selenite-F broth. A loopful of culture from Selenite-F was sub-cultured by streaking onto Xylose lysine deoxycholate agar (XLD) and incubated at 37 °C for 24h (Office International des Epizooties (OIE), 2012; Suresh et al., 2006). One millilitre of poultry drinking trough was inoculated into 9ml of BPW and the mixture was incubated at 37°C for 24h.

Thereafter, 1ml of the solution was transferred into 9ml Selenite-F broth and a loopful was streaked on XLD agar (Oxoid Basingstoke and Hampshire, UK) and incubated at 37 °C for 24 h.

### 2.2 Identification of presumptive isolates

Bacterial isolates were identified based on their cultural, biochemical, microscopic identification and morphological characteristics according to the method of Cheesbrough (Cheesbrough, 2006) and OIE (Office International des Epizooties (OIE), 2012). Isolates showing transparent colonies with black centre on XLD due to hydrogen sulphide production were selected as presumptive bacteria and subcultured on nutrient agar (Office International des Epizooties (OIE), 2012). Pure isolates were confirmed by their biochemical characteristics using urease, Triple sugar Iron test and citrate test. Afterwards, bacterial isolates identified as *Proteus* species were delineated into their various species by the indole test and stored on agar slants for further analysis.

### 2.3 Antimicrobial susceptibility and resistance profiling of *Proteus* species

The antimicrobial resistance profiles of *Proteus* species to ten antibiotics were determined by the disc diffusion method using standard procedures described by the Clinical Laboratory Standards Institute (CLSI) (CLSI, 2016). Ten commercial antibiotics disk (Oxoid, UK) which include: Amikacin (AMK) (30μg), Tetracycline (TET) (30μg), Ciprofloxacin (CPX) (5μg), Erythromycin (ERY) (15μg), Gentamycin (GEN) (10μg), Ampicillin (AMP) (10μg), Ofloxacin (OFX) (5μg), Ceftriaxone (CRO) (30μg), Levofloxacin (LVX) (5μg), and Penicillin (PEN) (10μg) were used.

Bacterial species were spread on Mueller Hinton agar and antibiotics discs were aseptically placed on the plates using sterile forceps. Plates were incubated for 24h at 37°C. Thereafter, the diameters of the zones of inhibition were measured in millimetre (mm) and results were interpreted using the Clinical Laboratory Standards Institute (CLSI) interpretative charts (CLSI, 2016).

### 2.4 Molecular Identification of Virulence Genes of *Proteus* species

#### 2.4.1 DNA extraction

Genomic DNA was extracted using the boiling method. Briefly, 5 mL of bacterial isolates grown in Laura Bertani (LB) broth at 37 °C for 8 h were centrifuged at 14000 rpm for 3 min. The cells were resuspended in 500 μl of normal saline and heated at 95 °C for 20 min in the heating chamber. The heated bacterial suspension was cooled on ice and centrifuged at 14000 rpm for 3 min. The supernatant containing the DNA was transferred to a 1.5 ml microcentrifuge tubes and stored at −20 °C for further use.

#### 2.4.2 DNA Quantification

The extracted genomic DNA was quantified using the NanoDrop 1000 spectrophotometer by placing a drop (approximately 2 μl) on the sample space and analysed using the NanoDrop 1000 software.

#### 2.4.3 Molecular characterisation of virulence genes of *Proteus* species

The virulence genes of *Proteus* species were characterised by Polymerase chain reaction technique. The following primer sets were used, (5’ – TTGAAGGACGCGATCAGACC – 3’) and (3’ – ACTCTGCTGTCCTGTGGGTA-5’) which amplifies a 467 bp sequence of *rsbA* gene (Abbas et al., 2015), (5’ – ACTCTGCTGTCCTGTGGGTA −3’) and (3’ – GTTATTCGTGATGGTATGGG-5’) which amplifies the 317 bp sequence of *ureC* gene (Pathirana et al., 2018) and (5’ – GTATGTCTGCACCTGCGGTA*-3’) and (3’ –* TTTGAGTTTGTCTTCTGGTAGTGC-5’) which amplifies the 464 bp sequence of *luxS* genes (Abbas et al., 2015). Gene amplification was carried out on thermal cycler (AB Biosystem, USA) at a final volume of 25 μl for 35 cycles. The PCR mix included X2 Dream Taq Master Mix supplied by Inqaba, South Africa (Taq polymerase, dNTPs, MgCl) and the primers at a concentration of 0.2M and 0.5 μl DNA as template. The PCR conditions were as follows: Initial denaturation, 95°C for 5 minutes; denaturation, 95°C for 30 seconds; annealing, 59°C for 30 seconds; extension, 72°C for 30 seconds for 35 cycles and final extension, 72°C for 5 minutes. The product was resolved on a 1% agarose gel at 120V for 20 minutes and visualised on a UV transilluminator.

## 3. Results

### 3.1 Identification of *Proteus* species

Presumptive isolates were identified based on their cultural and biochemical characteristics. Cultural characteristics included transparent colonies with black centre on XLD due to hydrogen sulphide production. A total of 74 isolates were confirmed positive on urease media with the production of pinkish-red colouration of the medium. Further examination using Triple sugar Iron test showed black butt from hydrogen sulphide production (H_2_S) while motility test produced a cloudy and distinct line of inoculation. Citrate test was positive for *Proteus* species. *P. vulgaris* was indole positive while *P. mirabilis* was indole negative.

The prevalence rate among the samples (n=216) of eggshells (88), poultry feed (n=64), drinking water (n=64) from the four poultry farms was 34.26% (74/216) (Table 1). About a quarter 23.86% (21/88) of eggshell swab samples were positive for *Proteus* while 39.06% (25/64) and 43.75% (28/64) of Feed and drinking water were positive for *Proteus* species respectively. Of the total 216 studied samples, Farm A and D had the highest prevalence rate of 9.26% (20/216), followed by Farm B and Farm C with prevalence rates of 8.80% (19/216) and 6.94% (15/216) respectively. However, the prevalence of *Proteus* sp. did not vary significantly (p ≤ 0.05) among all the samples collected from the four commercial farms.

**Table 1.** Prevalence of *Proteus* species from farm equipment and products from four commercial farms in Nasarawa State.

| Sample type | Prevalence of positive samples (%) |  |  |  |  |
| --- | --- | --- | --- | --- | --- |
|  | Farm A | Farm B | Farm C | Farm D | Total number of positive samples (%) |
| Eggshell swabs (n=22) | 5 (22.73) | 5 (22.73) | 4 (18.18) | 7 (31.82) | 21/88 (23.86) |
| Feed (n=16) | 7 (43.75) | 7 (43.75) | 5 (31.25) | 6 (37.5) | 25/64 (39.06) |
| Drinking water (n=16) | 8 (50.00) | 7 (43.75) | 6 (37.5) | 7 (43.75) | 28/64 (43.75) |
| Total (n=216) | 20 (9.26) | 19 (8.80) | 15 (6.94) | 20 (9.26) | 74 (34.26) |

From the sample types, water from the trough contained the highest amount of *Proteus* species 37.84% (28/74), followed by the feed 33.78% (25/74), then the eggshell 28.38% (21/74) as shown in Figure 1.

**Fig 1:**
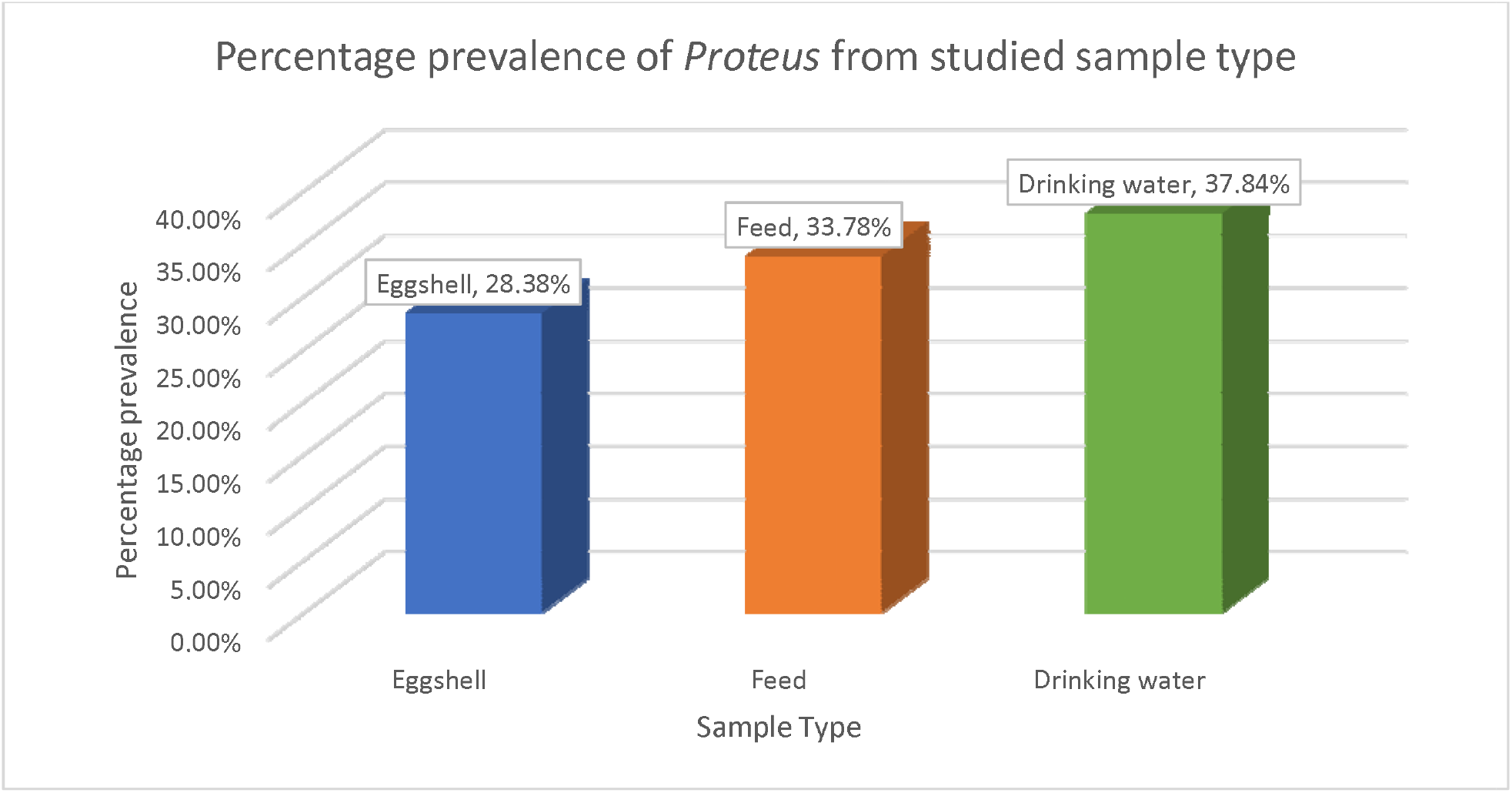
Percentage prevalence of *Proteus* species from the samples studied (n=74)

Table 2 shows that *P. mirabilis* was more prevalent 78% (58/74) than *P. vulgaris* 21.6% (16/74) among all the samples from the four farms. The drinking water sample contained the highest prevalence of *P. mirabilis* 82% (23/28), followed by feed sample 76% (19/25) and eggshell sample 76% (16/25) while the feed and eggshell samples had the highest prevalence of *P. vulgaris* 24% (6/25 and 5/21 respectively) and the lowest prevalence of 18% (5/28) was observed in drinking water samples. The prevalence of *P. mirabilis* was in the order; Farm A and C (80%), Farms B (79%) and Farm D (75%) while *P. vulgaris* was in the other Farm D (25%), followed by Farm B (21%) and farms A and C (20).

**Table 2.** Prevalence of *P. mirabilis* and *P. vulgaris* from four commercial farms in Nasarawa State. There was no significant statistical difference (p ≥0.05) between and within the *Proteus* species, farms and sample type. Key: ***Pm: Proteus mirabilis; Pv: Proteus vulgaris***

| Sample type | Farm A |  | Farm B |  | Farm C |  | Farm D |  | Cumulative Total across studied farms |  |
| --- | --- | --- | --- | --- | --- | --- | --- | --- | --- | --- |
|  | *Pm* (%) | *Pv* (%) | *Pm* (%) | *Pv* (%) | *Pm* (%) | *Pv* (%) | *Pm* (%) | *Pv* (%) | *Pm* (%) | *Pv* (%) |
| Egg shell | 4/5 (80) | 1/5 (20) | 4/5 (80) | 1/5 (20) | 3/4 (75) | 1/4 (25) | 5/7 (71) | 2/7 (29) | 16/21 (76) | 5/21 (24) |
| Feed | 5/7 (71) | 2/7 (29) | 6/7 (86) | 1/7 (14) | 4/5 (80) | 1/5 (20) | 4/6 (67) | 2/6 (33) | 19/25 (76) | 6/25 (24) |
| Drinking water | 7/8 (88) | 1/8 (12) | 5/7 (71) | 2/7 (29) | 5/6 (83) | 1/6 (17) | 6/7 (86) | 1/7 (14) | 23/28 (82) | 5/28 (18) |
| Total per *Proteus* sp | 16/20 (80) | 4/20 (20) | 15/19 (79) | 4/19 (21) | 12/15 (80) | 3/15 (20) | 15/20 (75) | 5/20 (25) | 58/74 (78) | 16/74 (22) |

### 3.2 Antimicrobial susceptibility and resistance profiling of *Proteus* species

As shown in Table 3, the antimicrobial susceptibility profile of the 74 confirmed *Proteus* species was tested against a panel of 10 antibiotics. *Proteus* species were susceptible to the antibiotics in varying amounts and susceptibility declined in the following order: Amikacin and Gentamicin (95.9%), Levofloxacin (93.2%), Ciprofloxacin and Penicillin (86.5%), Ofloxacin (82.4%), Erythromycin (81.1%), Ampicillin (77.0%) and Tetracycline (68.9%). The resistance of *Proteus* to antibiotics was observed in the following order with the highest resistance recorded for tetracycline (25.7%) followed by Erythromycin (13.5%), Cefotaxime (12.2%), Ampicillin (9.5%), Ofloxacin and Penicillin (5.4%), Gentamicin and Ciprofloxacin (4.1%) and Levofloxacin (1.4%). No resistance was observed for Amikacin.

**Table 3.** Antimicrobial susceptibility profile of *Proteus* species in the four commercial poultry farms in Nasarawa State.

| Antibiotics (µg) | Susceptible |  |  | Intermediate |  |  | Resistant |  |  |
| --- | --- | --- | --- | --- | --- | --- | --- | --- | --- |
|  | *P. mirabilis*<br>(%) | *P. vulgaris*<br>(%) | *Total* | *P. mirabilis*<br>(%) | *P. vulgaris*<br>(%) | *Total* | *P. mirabilis*<br>(%) | *P. vulgaris*<br>(%) | *Total* |
| Amikacin (30) | 55 (94.8) | 16 (100) | **71(95.9)** | 3 (5.2) | 0 (0) | **3 (4.1)** | 0 | 0 (0) | **0 (0)** |
| Ciprofloxacin (5) | 54 (93.1) | 10 (62.5) | **64(86.5)** | 3 (5.2) | 5 (31.3) | **8 (10.8)** | 0 (0) | 2 (12.5) | **3 (4.1)** |
| Erythromycin (15) | 51 (87.9) | 9 (56.3) | **60(81.1)** | 2 (3.4) | 2 (12.5) | **4 (5.4)** | 5 (8.6) | 5 (31.3) | **10 (13.5)** |
| Tetracycline (30) | 45 (77.6) | 6 (37.5) | **51(68.9)** | 4 (6.9) | 0 (0) | **4 (5.4)** | 9 (15.5) | 10 (62.5) | **19 (25.7)** |
| Levofloxacin (5) | 53 (91.4) | 16 (100) | **69(93.2)** | 4 (6.9) | 1 (6.3) | **5 (6.8)** | 1 (1.7) | 0 (0) | **1 (1.4)** |
| Ofloxacin (5) | 53 (91.4) | 8 (50) | **61(82.4)** | 3 (5.2) | 6 (37.5) | **9 (12.2)** | 3 (5.2) | 1 (6.3) | **4 (5.4)** |
| Cefotaxime (30) | 48 (82.3) | 11 (68.8) | **59(79.7)** | 4 (6.9) | 2 (12.5) | **6 (8.1)** | 6 (10.3) | 3 (18.8) | **9 (12.2)** |
| Gentamicin (10) | 57 (98.3) | 14 (87.5) | **71(95.9)** | 0 (0) | 0 (0) | **0 (0)** | 2 (3.4) | 1 (6.3) | **3 (4.1)** |
| Ampicillin (10) | 49 (84.5) | 8 (50) | **57(77.0)** | 6 (10.3) | 4 (25.0) | **10 (13.5)** | 3 (5.2) | 4 (25.0) | **7 (9.5)** |
| Penicillin (10) | 51 (87.9) | 13 (81.3) | **64(86.5)** | 5 (8.6) | 1 (1.4) | **6 (8.1)** | 2 (3.4) | 2 (12.5) | **4 (5.4)** |

### 3.3 Molecular characterisation of virulence genes of bacterial species

The molecular analysis of *Proteus* species was carried out to detect the virulence genes *luxS, ureC* and *rsbA* genes. Among the 15 *Proteus* species, the *luxS* gene was detected in seven *Proteus* isolates (four isolates of *P. vulgaris* and three isolates of *P. mirabilis*) (Figure 2). The *ureC* gene was detected in ten Proteus isolates (six isolates of *P. mirabilis* isolates and four *P. vulgaris* isolates) (Figure 3) while the *rsbA* gene was detected in four *Proteus* isolates (three *P mirabilis* and one *P. vulgaris* isolates) (Figure 4).

**Figure 2:**
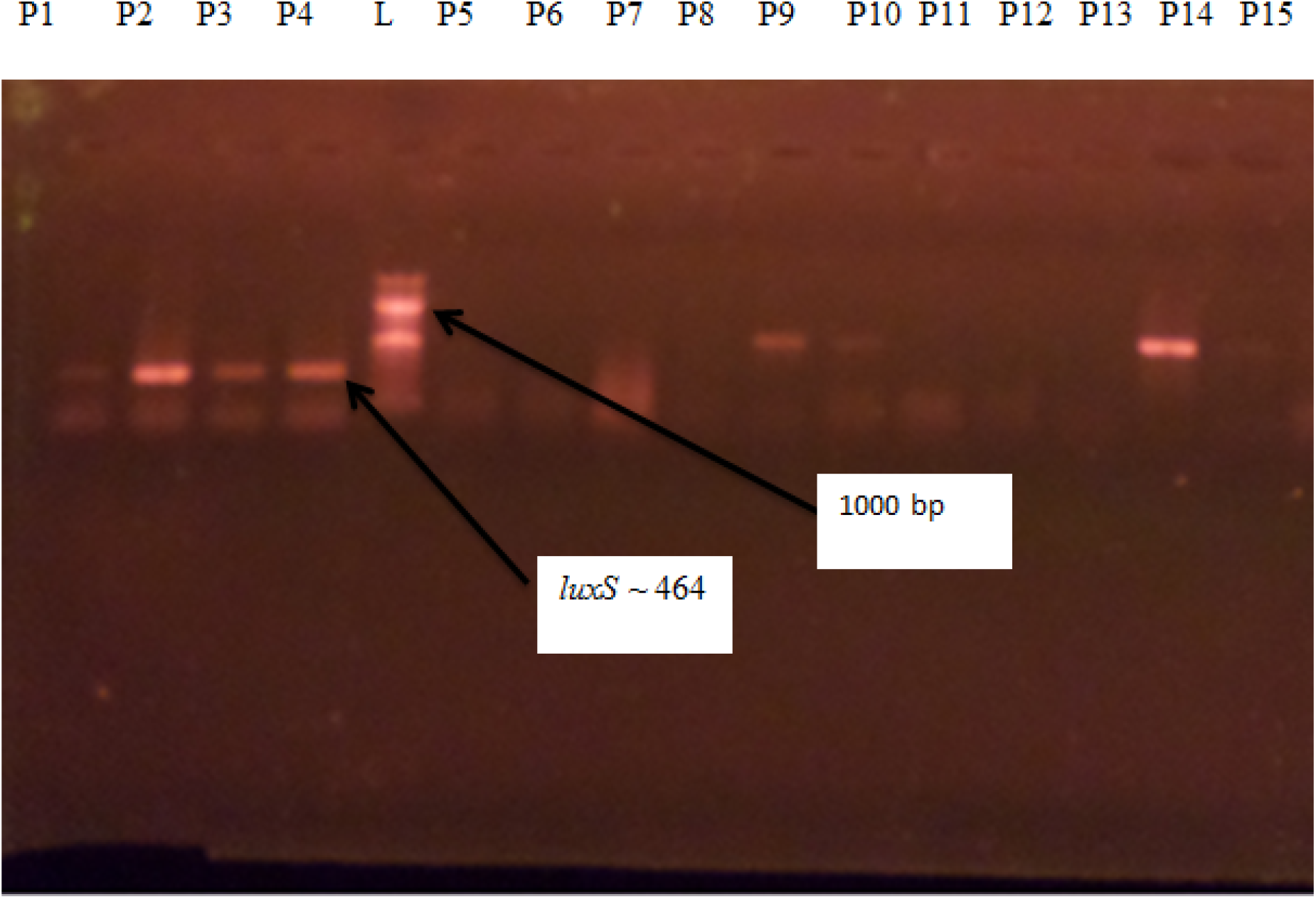
Agarose gel electrophoresis of the amplified *luxS* genes from the *Proteus* species. Lanes P1-P4, P9, P10 and P14 represent the *luxS* bands. Lane L represents the 1000 bp molecular ladder **Key**: P1-P5 represent *P. vulgaris;* P6-P15 represent *P. mirabilis.*

**Figure 3:**
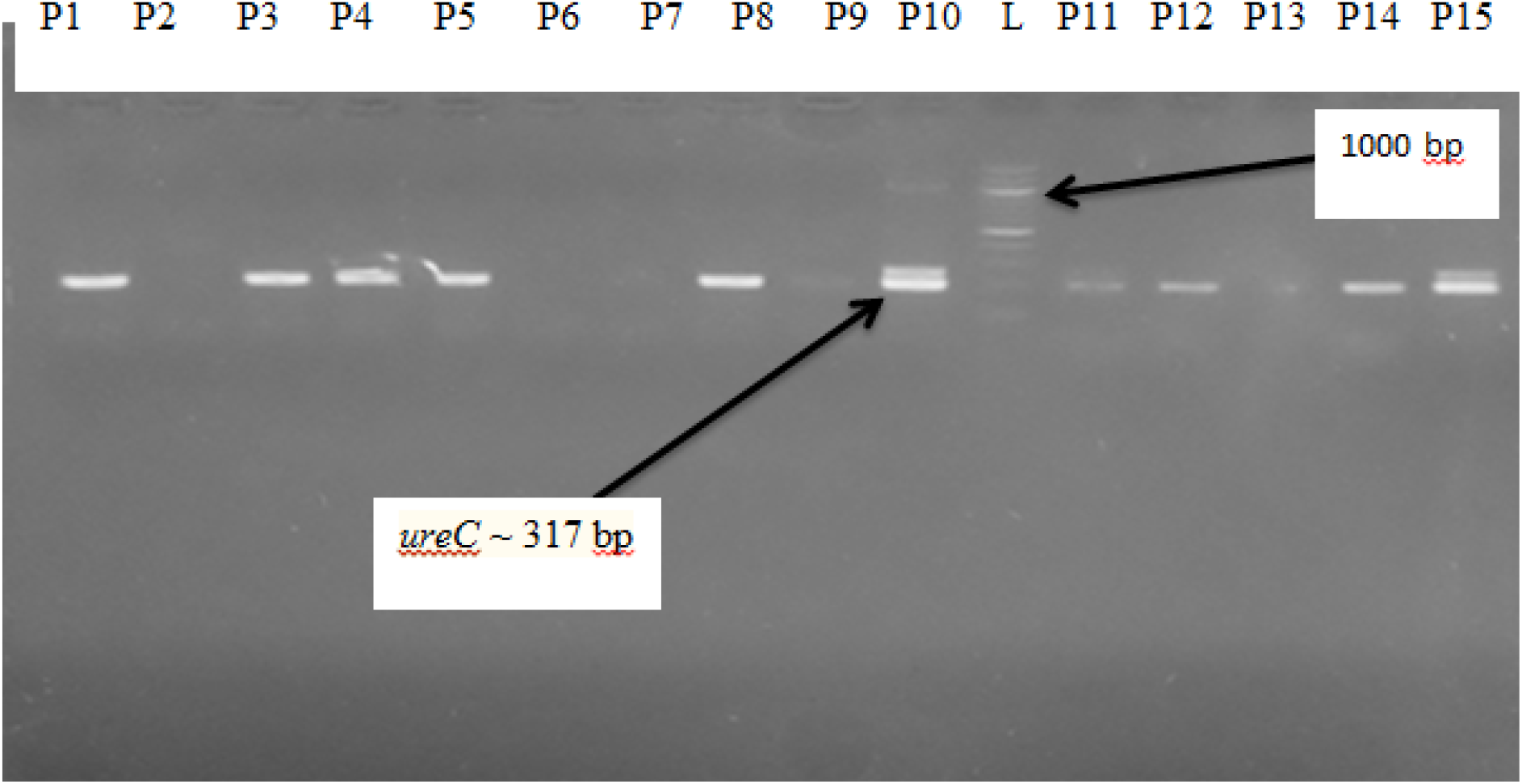
Agarose gel electrophoresis of the amplified *ureC* genes from the *Proteus* species. Lanes P1, P3-P5, P8, P10, P11, P12, P14 and P15 represent the *ureC* bands. Lane L represents the 1000 bp molecular ladder, while other lanes show no band. **Key**: P1-P5 represent *P. vulgaris;* P6-P15 represent *P. mirabilis.*

**Figure 4:**
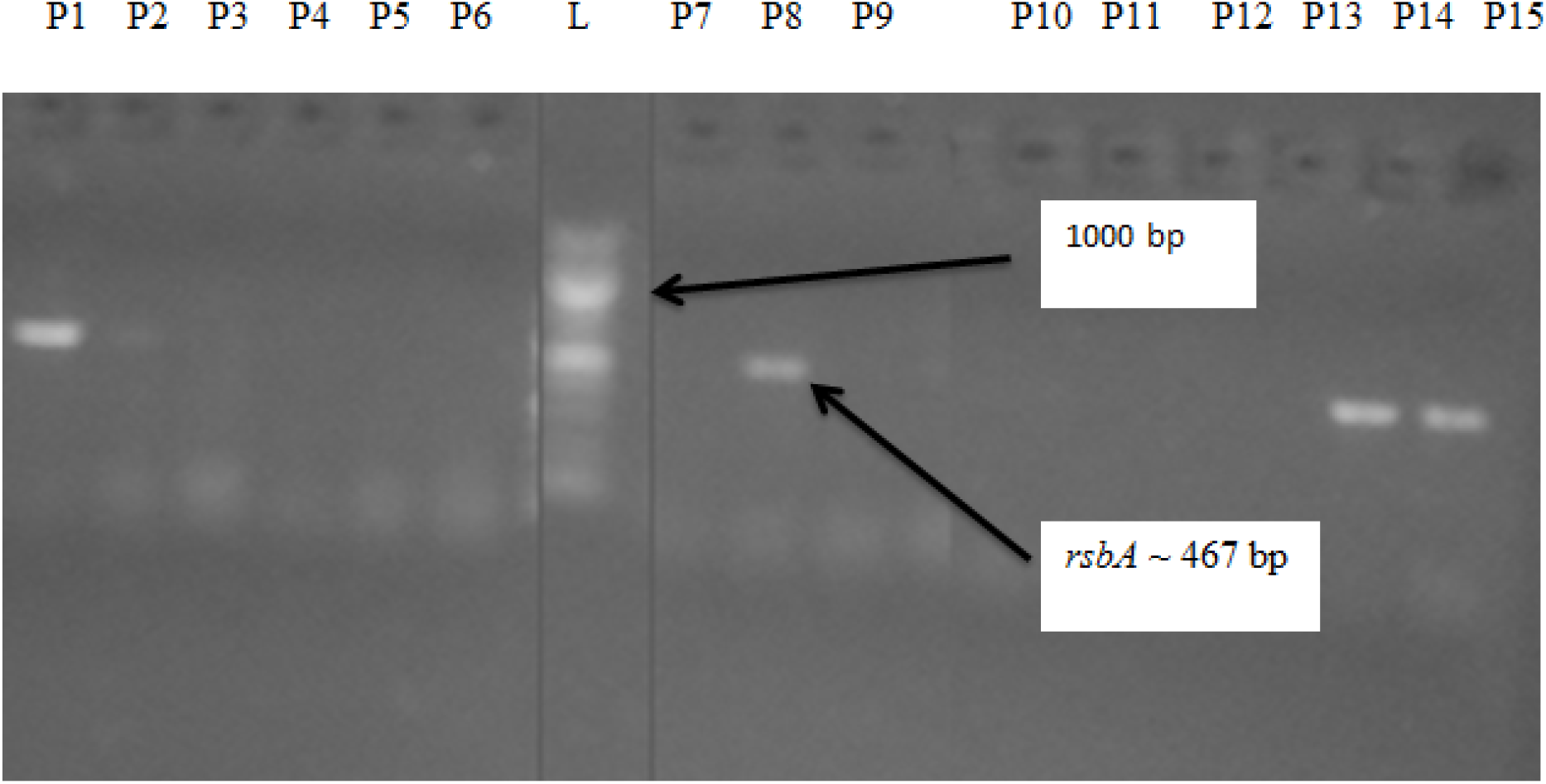
Agarose gel electrophoresis of the amplified *rsbA* genes from the *Proteus* species. Lanes P1, P8, P14 and P15 represent the *rsbA* bands. Lane L represents the 1000 bp molecular ladder, while other lanes showed no band. **Key**: P1-P5 represent *P. vulgaris;* P6-P15 represent *P. mirabilis.*

## 4. DISCUSSION

The cross-sectional study examined the phenotypic resistance and virulence profile of *Proteus* species from eggshells, feed and drinking water samples collected from four commercial poultry farms in Lafia, Nasarawa State. The overall prevalence of *Proteus* species was approximately 34.3% (74 positive samples, n= 216) with drinking water having the highest number of positive samples and *P. mirabilis* was the predominant species found in all the samples. Although the prevalence of *Proteus* species among all the samples from the four farms was not statistically significantly different (P ≤ 0.05), the number of isolated *Proteus* species is still worrisome due to the ability of these bacterial pathogens to cause human infections such as pneumonia, septicaemia, central nervous system infection, food poisoning and urolithiasis (Barbour et al., 2012). Direct contact of humans with faecal droppings harbouring *Proteus* species can result in the transmission of the pathogens to immune-suppressed persons leading to urinary tract infections (Pathirana et al., 2018). *P. mirabilis* has been frequently isolated from chicken eggs, cloacal swabs and environmental samples from poultry farms. Ubiebi (Ubiebi, 2017) isolated *P. mirabilis* from poultry feeds in Delta State, Nigeria and *P. mirabilis* were among the isolates present in layers mash and broiler finisher samples obtained from ten poultry farms in Ekiti, Nigeria (Oyinloye et al., 2015). Enterobacteriaceae, including *P. mirabilis* and *P. vulgaris,* were frequently isolated from poultry birds (Kuznetsova et al., 2019).

Data generated from this study is of great public health importance as empirical evidence has demonstrated Proteus species’ involvement in human diseases and as agents of animal infections and bacterial contamination of poultry products (Nahar et al., 2014). Additionally, the occurrence of *Proteus* species in these farms indicates defective biosafety measures in controlling rodents’ infestations, insect vectors, wild birds and pet movements in the poultry pens.

The high prevalence of *Proteus mirabilis* (78%) reported in this study is comparably higher to the prevalence of 66% documented by (Barbour et al., 2012). The highest prevalence of *Proteus* species, notably *P. mirabilis*, was recorded for farms A and C while Farm D had the lowest Proteus species’ prevalence. This result could be attributed to poor sanitary practices in the farms such as lack of control of the entry of rodents and wild birds into the poultry houses, poor cleaning and disinfection of pens, and the citing of the two farms in waterlogged areas. On the other hand, Farm D was well fenced and good hygienic practices such as frequent cleaning of feed and water troughs were observed. Additionally, the entry of visitors into farm D is restricted in contrast to unrestricted visits to Farms A-C.

Antibiotics resistant bacteria were detected in all the four farm samples. The 25.7% resistance to Tetracycline in this study disagrees with the result of (Dadheech et al., 2015) who recorded a 100% resistance of *Proteus* species isolated from chicken to Tetracycline in Ajmer Region, India. Data obtained may be attributed to tetracycline’s frequent use in routine prophylaxis and chemotherapy in livestock management in Nigeria (Aliyu et al., 2019). The isolates also exhibited varying resistance to the other antibiotics such as Erythromycin (13.5%) and Cefotaxime (12.2%). This finding is baffling considering that Erythromycin and Cefotaxime are not routinely used in poultry management in the areas studied. This finding may suggest the transmission of Erythromycin and Cefotaxime resistant strains to the birds or contamination of eggs/water by poultry attendants in the farms. Antibiotics may be continuously administered as growth promoters to food-grade animals such as broilers and turkeys; therefore, the antibiotic selection pressure for resistance amongst bacteria in poultry is high and consequently their faecal flora contains a relatively high proportion of resistant bacteria (Nair et al., 2017). The interaction between the different components in a food chain or the environment further contributes to the spread of antibiotic resistance across species (Landers et al., 2012).

The most effective antibiotics against *Proteus* species were the Aminoglycosides, Amikacin and Gentamicin, with 95.9% activity against *Proteus* while the highest resistance was recorded for tetracycline (25.7%). Similar findings reported by (Kuznetsova et al., 2019) showed a low resistance of *Proteus* to Amikacin and Gentamicin and total resistance to tetracycline. The resistance of *Proteus* species to Ciprofloxacin (4.1%) was low in this study compared to the report of Kuznetsova *et al.* (Kuznetsova et al., 2019) who recorded a 36.8% resistance to Ciprofloxacin. The low level of resistance exhibited by the isolates to Ciprofloxacin may be attributed to the small size of the drug which is a factor that enhances its solubility in the diluents thus enhancing its penetration power through the cell into the cytoplasm of the target organism where it exerts its effects (Okpo et al., 2017). The high resistance of multidrug *P. mirabilis* to Tetracycline and its susceptibility to Ciprofloxacin in droppings of poultry birds collected in Bangladesh (Nahar et al., 2014) is consonance with the result obtained in our study where *P. mirabilis* was resistant to Tetracycline and sensitive to Ciprofloxacin. Okonko *et al.* (Okonko et al., 2010) found *P. mirabilis* to be 100% sensitive to streptomycin, an aminoglycoside in the same family with Amikacin and Gentamicin used in our study where the sensitivity was 95.9%. Likewise, the resistant and susceptibility patterns of *P. mirabilis* to tetracycline and Gentamicin respectively were established in poultry feeds purchased from Calabar, Nigeria (Okonko et al., 2010). This report suggests that the presence of *Proteus* in poultry feeds may be attributed to commercially sold feeds contaminated with *Proteus* and not necessarily from poultry workers directly infecting the feed. In agreement with similar studies, the aminoglycosides seem to be the drug of choice to treat *Proteus* infections. The extra outer cytoplasmic membrane comprising the lipid bilayer, lipoproteins, polysaccharides and lipopolysaccharides may be responsible for *Proteus* resistance to antibiotics (Okonko et al., 2010). The availability and accessibility to antimicrobial compounds such as Ciprofloxacin, Streptomycin, Gentamicin, Erythromycin, Tetracycline, and Furazolidone among others in open markets to treat broiler/layer pose a challenge to reducing antimicrobial resistance in the poultry farms (Hasan et al., 2011; Nahar et al., 2014).

The *ureC* gene was the predominant gene detected in this study. *ureC* gene is responsible for the elevation of urine pH, resulting in stone formation and it plays a crucial role in the virulence of *Proteus* species (Mohammed et al., 2014). Ram *et al.* (Ram et al., 2019) found *ureC* genes in the chicken cloacal swabs collected from a livestock farm complex in India. The frequency of occurrence of *ureC* genes (66.7%) among the *Proteus* species was higher than the other two genes. However, the frequency of 66.7% is lower than similar findings of Pathirana *et al.* (Pathirana et al., 2018) who isolated *Proteus* species from humans and pet turtles and reported 91.7% prevalence for *ureC* genes. Gene expression of *ureC* genes and other virulent genes in Multidrug-resistant *P. mirabilis* isolated from diarrhetic animals in North-East China were associated with biofilm formation (Sun et al., 2020). The *rsbA* gene was amplified in both *P. mirabilis and P. vulgaris* with a 26.7% frequency which is in contrast to the report of Abbas *et al.* (Abbas et al., 2015), who reported that *rsbA* gene could not be amplified in *P. mirabilis and P. vulgaris*. Swarming behaviour of *P. mirabilis* is mediated by *rsbA* gene, which may function as a protein sensor of environmental conditions (Różalski et al., 2012). The *rsb*A gene is also responsible for biofilm formation and extracellular polysaccharide formation (Różalski et al., 2012). The *lux*S genes were detected in all the *Proteus* species. *luxS* genes production has been implicated in biofilm formation by members of the Enterobacteriaceae, including *Proteus* species (Badi and Sepahi, 2014).

## 5. CONCLUSIONS

This study revealed the prevalence of antimicrobial-resistant *Proteus* species in poultry feed, water and eggshells collected from four poultry farms in Lafia, Nigeria. *Proteus* species were also confirmed to possess virulence genes suggestive of potential threat to food safety. To minimise the spread of antimicrobial resistance arising from poultry farms, it is vital to regulate the use of antimicrobial agents as growth promoters for poultry birds. Effective management of the poultry farm environment and farm products can also be achieved through routine monitoring and surveillance of farms to avert possible transmission of pathogenic microorganisms via the food chain. It is also a matter of urgency to ensure proper sanitation and hygiene in the farms by farm managers. Quality control of feeds purchased should be carried out to avoid transmission of pathogens from external sources. These measures will enable the check of pathogenic bacteria, including *Proteus* from poultry farms.

## Acknowledgement

The authors hereby acknowledge Dr. Aliyu Yakubu (Department of Science Laboratory Technology, Federal Polytechnic Nasarawa) and the Department of Microbiology Laboratory’s support staff, Federal University of Lafia for their support during the conduct of this study.

## Conflict of interest

The authors declare that there is no conflict of interest.

## Funding source

This research did not receive any specific grant from funding agencies in the public, commercial, or not-for-profit sectors.

## Notes

### Competing Interest Statement

The authors have declared no competing interest.

### Summary of Updates

To correct typographical, data and statistical errors.

